# The SWI/SNF nucleosome remodeler constrains enhancer activity during *Drosophila* wing development

**DOI:** 10.1101/2023.07.17.549384

**Authors:** Matthew J. Niederhuber, Mary Leatham-Jensen, Daniel J. McKay

**Affiliations:** Curriculum in Genetics and Molecular Biology, The University of North Carolina at Chapel Hill, Chapel Hill, NC, 27599; Department of Biology, The University of North Carolina at Chapel Hill, Chapel Hill, NC, 27599; Department of Genetics, The University of North Carolina at Chapel Hill, Chapel Hill, NC, 27599; Integrative Program for Biological and Genome Sciences, The University of North Carolina at Chapel Hill, Chapel Hill, NC, 27599

**Keywords:** nucleosome remodeler, chromatin, enhancer, developmental gene regulation

## Abstract

Chromatin remodeling is central to the dynamic changes in gene expression that drive cell fate determination. During development, the sets of enhancers that are accessible for use change globally as cells transition between stages. While transcription factors and nucleosome remodeling complexes are known to work together to control access to enhancers, it is unclear how the short stretches of DNA that they individually unmask yield the kilobase-sized accessible regions that are characteristic of active enhancers. Here, we performed a genetic screen to investigate the role of ATP-dependent nucleosome remodeling complexes in the control of dynamic enhancer activity. We find that the *Drosophila* BAP complex, a member of the SWI/SNF family of nucleosome remodelers, is required for repression of a temporally dynamic enhancer, *br^disc^*. Contrary to expectations, we find that the BAP-specific subunit, Osa, is dispensable for mediating changes in chromatin accessibility between early and late stages of wing development. Instead, we find that Osa is required to constrain the levels of *br^disc^* activity in imaginal wing discs when the enhancer is normally active. Genome-wide profiling reveals that Osa binds directly to the *br^disc^* enhancer as well as thousands of other developmentally dynamic regulatory sites, including multiple genes encoding components and targets of the Notch signaling pathway. We find that Osa loss of function results in development of ectopic sensory structures that are normally patterned by Notch signaling early in wing development. Moreover, we find that Osa loss of function results in hyperactivation of the *Delta* gene, which encodes the Notch ligand. Together, these findings indicate that proper constraint of enhancer activity is necessary for regulation of dose-dependent developmental events.

## INTRODUCTION

Animal development requires robust control over the spatial patterns, magnitude, and temporal dynamics of gene expression. Dysregulation in any of these regulatory dimensions is known to contribute to developmental disorders and acquired disease states. Spatial control refers to the selective patterns of gene expression across a field of cells. For instance, the spatially restricted expression of Hox genes in animals is essential for specification of regional identities along the developing body axis (Mallo *et al*. 2013). Both loss of expression and ectopic expression of Hox genes beyond their normal spatial domains can lead to homeotic transformations. The magnitude of gene expression must also be tightly controlled for proper development, and both excessive and insufficient gene expression can be detrimental. For instance, duplication of the *APP* gene is associated with early onset Alzheimer’s disease and is thought to be a driver of Alzheimer’s in individuals with Trisomy 21 (Tang *et al*. 2013). Conversely, heterozygosity of Notch pathway components, including the Notch receptor itself, is associated with several developmental syndromes (Falo-Sanjuan and Bray 2020). This dose-dependency is conserved in *Drosophila*, which exhibit defects in sensory organ development when genes encoding Notch pathway components are mutated, as well as in genotypes with extra copies of Notch pathway genes (Hartenstein and Posakony 1990; Parks and Muskavitch 1993; Doherty *et al*. 1996; Elfring *et al*. 1998; Armstrong *et al*. 2005). The spatial patterns and levels of gene expression are also temporally dynamic, as cells transition through intermediate identities over developmental time. A classic example of temporal regulation is ecdysone hormone signaling in insects, which triggers changes in stage-specific gene expression programs across body parts that are not in close physical contact (Yamanaka *et al*. 2013). Despite their importance, the factors and mechanisms coordinating these three dimensions of developmental gene regulation remain incompletely understood.

A primary layer of gene regulation lies at the level of *cis*-acting DNA regulatory elements and the *trans*-acting factors that bind them. Enhancers are relatively short (∼0.5-2kb) non-coding regions of DNA that function as integration points for the spatiotemporal information transmitted by sequence-specific transcription factors, which typically bind short DNA sequences 6-10bp in length (Spitz and Furlong 2012; Uyehara and Apostolou 2023). Additional layers of information come in the form of the packaging and chemical modification of chromatin. Histone post-translational modifications directly and indirectly control chromatin structure and help propagate cellular memory (Millán-Zambrano *et al*. 2022). Access to DNA-encoded information is also influenced by the positioning, stability, and occupancy of nucleosomes. Nucleosomes are inhibitory to transcription factor binding and thus must be remodeled or removed for an enhancer to become active (Brahma and Henikoff 2020; Niederhuber and McKay 2020; Isbel *et al*. 2022). Genome-wide patterns of chromatin accessibility are predictive of enhancer activity. Moreover, temporal changes in chromatin accessibility profiles are correlated with stage-specific changes in gene expression during development (Uyehara *et al*. 2017). Recent studies in *Drosophila* have provided insight into the mechanisms controlling developmentally programmed changes in chromatin accessibility. A number of transcription factors have been identified that open chromatin in early stage embryos to promote activation of the zygotic genome (Gaskill *et al*. 2021). Likewise, the ecdysone-induced transcription factor E93 has been found to be required for promoting accessibility of enhancers active later in pupal stages of development (Nystrom *et al*. 2020). Interestingly, E93 is also necessary for closing and deactivating early acting enhancers (Uyehara *et al*. 2017; Nystrom *et al*. 2020). Returning accessible enhancers to a closed chromatin state is important for rendering them refractory to transcription factor binding, thereby allowing regulatory inputs to be utilized at distinct targets over the course of development. However, the mechanisms of closing chromatin to repress enhancers during development are poorly understood relative to those controlling chromatin opening.

Here, we examine the contribution of nucleosome remodelers in control of a developmentally dynamic enhancer in *Drosophila*. Nucleosome remodelers use ATP hydrolysis to disrupt histone-DNA interactions, and by doing so, occlude or make accessible short stretches of DNA to transcription factors. Through mechanisms that remain unclear, disruption of short stretches of histone-DNA contacts by nucleosome remodelers can result in accessibility of enhancers that are often orders of magnitude greater in length (Clapier *et al*. 2017). Through an *in vivo* RNAi screen, we identified the *Drosophila* BAP complex, which is orthologous to yeast and human SWI/SNF, as being required for repression of a developmentally dynamic enhancer. Contrary to expectations, we find that BAP is dispensable for developmentally programmed changes in chromatin accessibility during wing metamorphosis. Instead, we find that the BAP subunit Osa is required to constrain activity when the enhancer is in the on state. Using CUT&RUN, we find that Osa directly binds thousands of regions that have signatures of active enhancers, including multiple genes in the Notch signaling pathway. Lastly, we find that loss of BAP function results in upregulation of a direct Osa target gene, *Delta*, which encodes the Notch ligand. Together these data suggest a model in which the BAP complex directly constrains enhancer activity to ensure correctly measured responses to developmental signals like Notch signaling and cell specification programs during wing development.

## RESULTS

### The *br^disc^* enhancer is a model of a developmentally dynamic regulatory element

In order to interrogate the role of nucleosome remodelers in developmentally dynamic enhancer regulation, we selected a previously identified enhancer known to respond to temporal inputs from the ecdysone hormone pathway (Uyehara *et al*. 2017; Nystrom *et al*. 2020). The *br^disc^* enhancer is a ∼2kb region on the X chromosome that lies approximately 9kb upstream of the gene *broad* (*br*). Prior studies of *br^disc^* activity using transgenic reporters indicated that it switches on prior to the 3^rd^ larval instar stage in the precursors of the adult appendages, including the wing (Uyehara *et al*. 2017). *Br^disc^* is then deactivated during the first 24 hours of pupal development, thus making it a good model for studying temporally dynamic enhancer control.

We first sought to improve the temporal resolution of *br^disc^*transgenic reporter activity. Traditional enhancer reporters optimize rapid fluorophore maturation, brightness, and stability. Although these optimizations are useful for sensitive detection of enhancer activity patterns, they are problematic when monitoring dynamic enhancer behavior because persistent fluorescent protein interferes with determining when enhancer activity shuts off. To mitigate these effects, we developed two new fluorescent reporters. The first is a dual fluorophore reporter system, which we refer to as *br^disc^-switch*. This reporter was designed to drive expression of a tandem tomato fluorophore (tdTomato) that could be inducibly switched via FLP/FRT-meditated recombination to transcribe a myristoylated GFP (myr-GFP, **Fig 1A**). The *switch* reporter system allows for better temporal resolution of enhancer dynamics relative to conventional reporters because GFP detected after reporter switching indicates the enhancer was active after recombination was induced. Conversely, a lack of GFP detected after reporter switching demonstrates that the enhancer was inactive at the time of recombination or later. Examination of *br^disc^-switch* activity revealed that the reporter is highly active in third larval wandering (3LW) wing imaginal discs, but there is little to no detectable nascent GFP in young pupal wings aged <40 hours after puparium formation (APF) (**Fig 1B**). By contrast, tdTomato signal remains high in this same wing (**Fig 1B**). Interestingly, the *br^disc^-switch* reporter exhibits new GFP signal in older (>40h APF) wings, in bristle shafts located along the wing margin and in cells of the posterior cross vein (PCV), indicating that it is reactivated in a subset of pupal wing cells after its initial deactivation. Closer inspection of another transgenic *br^disc^* reporter integrated at a separate genomic locus, and which employs a different minimal promoter, revealed similar late enhancer activity along the wing margin (**Fig S1A**). Thus, the observed spatiotemporal changes in reporter expression are likely driven by the enhancer rather than DNA sequences in the vector or surrounding genomic regions.

**Figure 1.**
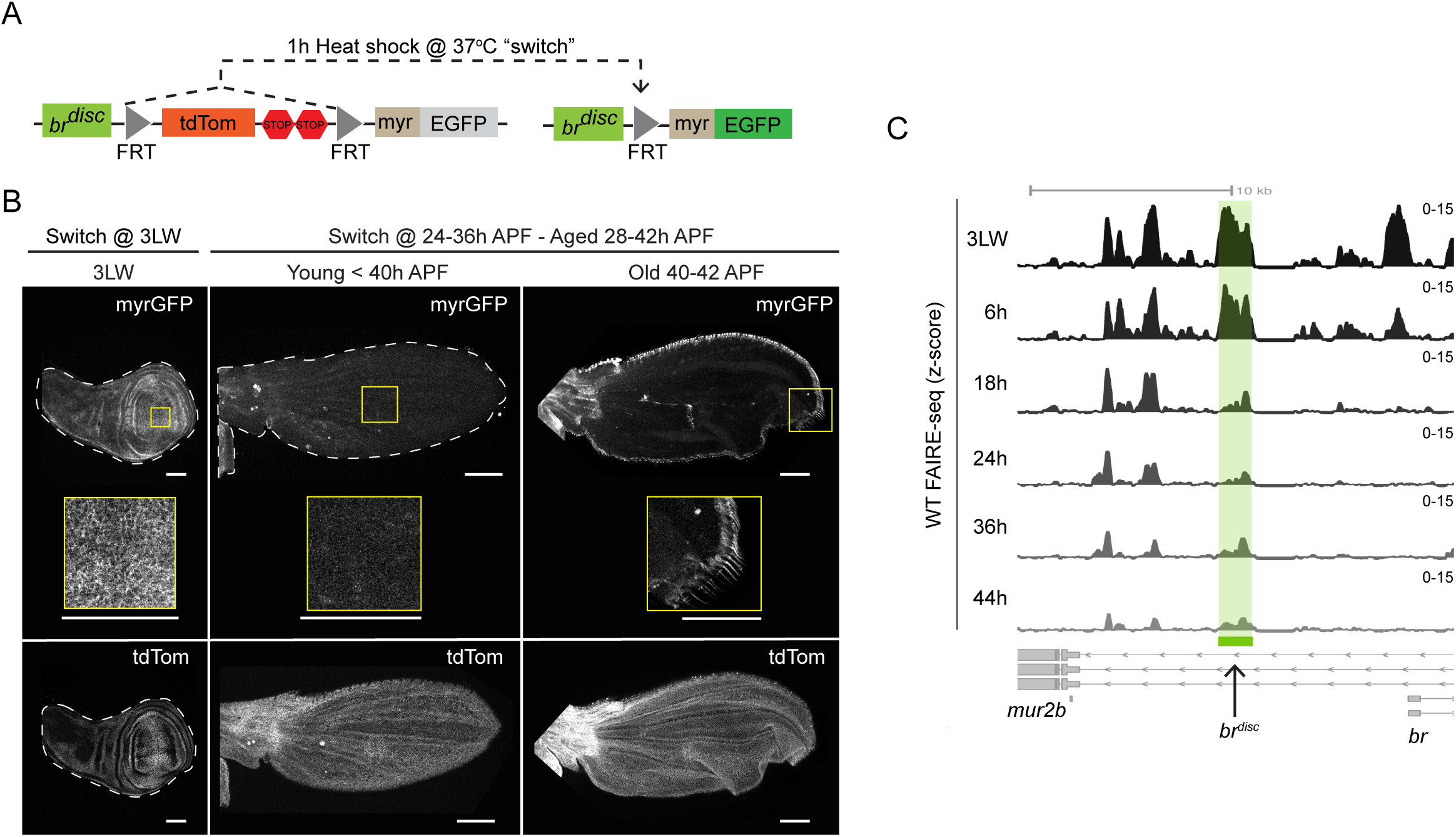
The *br^disc^*enhancer is a model of a developmentally dynamic regulatory element. (A) Illustration of the *br^disc^-switch* reporter. Heat shock-induced FLP expression excises the FRT-flanked “*tdTomato, 2xSTOP”* cassette to allow expression of myr-GFP. (B) Confocal images of *br^disc^-switch* activity in 3^rd^ Larval Wandering (3LW) imaginal wing discs and pupal wings aged 28-42h APF. “Young” and “Old” denote pupal wings categorized by morphology (see Methods). “Switch @” denotes ages of animals at time of heat shock. Images pupal wings are maximum intensity projections. Image of imaginal wing disc is single slice. Scale bars are 100µm. Wings are shown with anterior up. (C) Genome browser shot of z-normalized FAIRE-seq signal at the *br^disc^* enhancer (green highlight) from time course of WT wing development.

Comparison of temporal changes in *br^disc^* reporter activity and a chromatin accessibility time course performed in developing wild-type wings revealed a strong correlation between reporter activity and endogenous enhancer accessibility. *Br^disc^* exhibits high accessibility in 3LW wing discs, remains in an open state during the prepupal stage at 6h APF, and subsequently loses most of its accessibility by 18h APF, shortly after the prepupal to pupal transition (Uyehara *et al*. 2017; **Fig 1C**). These accessibility profiles are congruent with changes in reporter activity. We note that the later reactivation of *br^disc^* along the pupal wing margin does not coincide with a detectable increase in accessibility. This may be due to a lack of sensitivity to detect changes in a small number of cells using whole-wing FAIRE-seq. Alternatively, the small amount of accessibility that remains at later stages may derive from this population of cells. Together these observations demonstrate that *br^disc^* is dynamically active and accessible during wing development, thus making it a useful model for studying the mechanisms of dynamic enhancer regulation.

### The *Drosophila* BAP nucleosome remodeling complex is required to repress *br^disc^*

To identify factors that contribute to the developmental dynamics of *br^disc^* enhancer activity during wing metamorphosis we performed an *in vivo* RNAi screen. As described above, conventional fluorescent reporters are engineered to be highly stable, making them poorly suited for detecting changes in enhancer activity. While the *switch* reporter corrects for this problem by using a two-fluorophore output, it is too technically cumbersome for use in a larger scale screen. To circumvent this limitation while optimizing screen throughput, we created a second new transgenic fluorescent reporter in which the *br^disc^*enhancer drives tdTomato fused to a C-terminal PEST degradation tag (*br^disc^-tdT-PEST*; Li *et al*. 1998; Nern *et al*. 2011). This design yielded increased sensitivity for detecting *br^disc^* dysregulation, as determined by comparing reporter levels in the presence and absence of the PEST tag upon knockdown of a known *br^disc^* repressor (**Fig S2A**). We interpret the increase in sensitivity to be due to increased tdTomato turnover relative to the non-tagged version. Despite addition of the PEST tag, a low but detectable level of tdTomato expression was observed in wild-type pupal wing cells, which we interpret as residual fluorophore expression from earlier times in development when the enhancer is active (**Fig S2A**). RNAi expression was controlled by the UAS/GAL4 system via the *cubitus interruptus* anterior compartment GAL4 driver (*ci-GAL4*). A ubiquitously expressed temperature-sensitive allele of the GAL4 repressor GAL80 (*tub-GAL80^ts^*) was used to restrict RNAi expression to later stages of development (*ci^ts^*; **Fig 2A**). We envisioned two potential outcomes upon RNAi knockdown of *br^disc^* regulators: loss of an activator would yield decreased levels of tdTomato relative to control cells, and loss of a repressor would yield increased levels of tdTomato relative to control cells. We reasoned that by screening for tdTomato levels in pupal wings, we would potentially capture two types of repressors, those that deactivate *br^disc^* over time and those that constrain the levels of *br^disc^* activity while it is on in larval stages (**Fig 2B**). Knockdown of the transcription factor Eip93F (E93), a known negative regulator of *br^disc^*expressed during pupal stages, resulted in increased *br^disc^* reporter activity in pupal wings, whereas expression of a negative control *lexA*-RNAi failed to impact *br^disc^* activity, confirming the sensitivity of our reporter screen design to detect changes in enhancer activity (**Fig 2C**).

**Figure 2.**
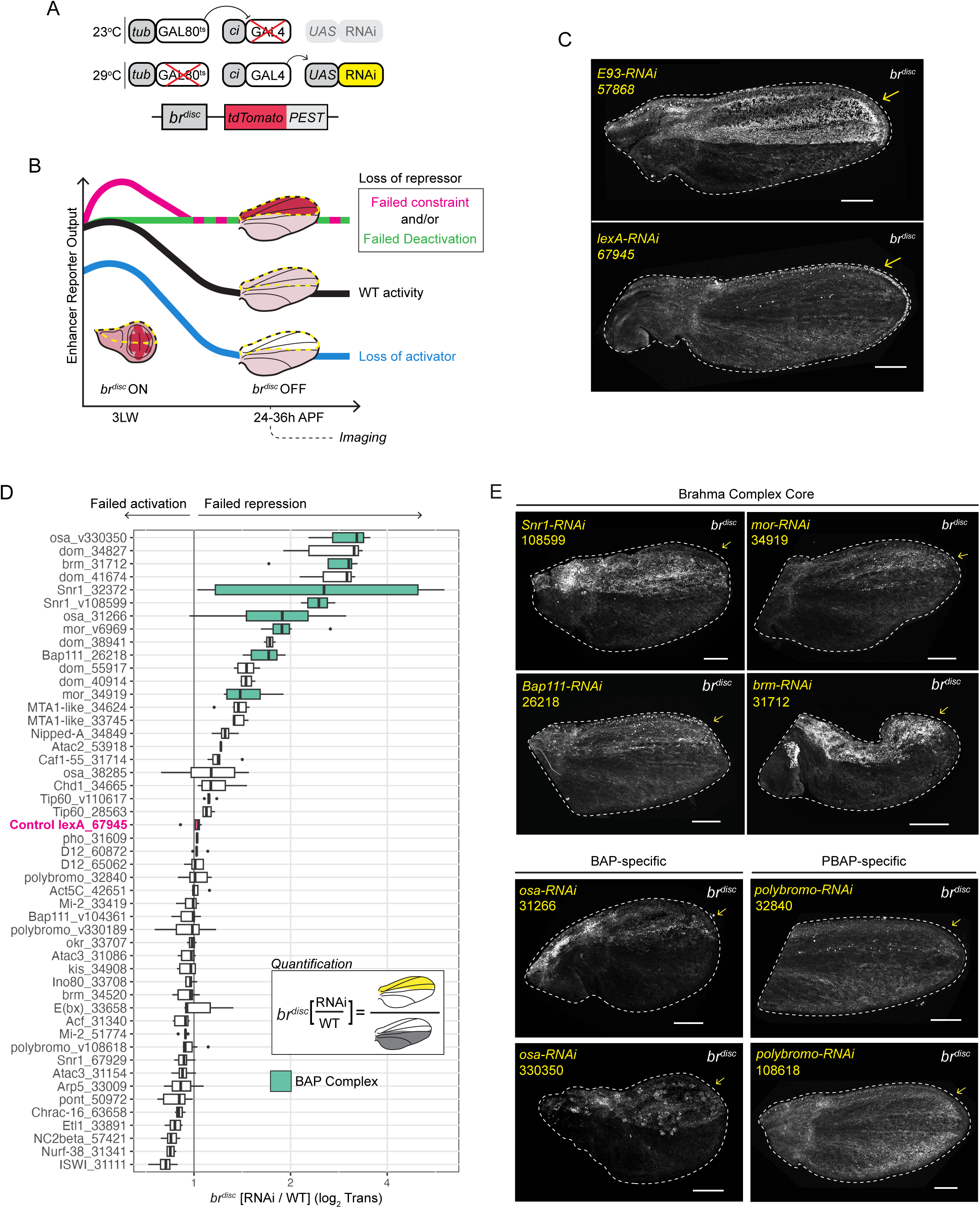
The BAP Complex is required to repress the *br^disc^* enhancer. (A) Illustration of the *br^disc^-tdTomato-PEST* reporter and inducible RNAi system used to screen for genes required for *br^disc^* regulation. (B) Schematic of types of enhancer dysregulation detectable in the RNAi screen. RNAi-expressing cells are located within the yellow dashed outline. Temporal activity of reporter in WT cells is indicated by black line. Loss of an activator in 3LW imaginal wing discs (*br^disc^* ON) would cause reduced reporter levels in RNAi cells (Blue line). Failed deactivation (*br^disc^*OFF) would cause increased levels of reporter activity in RNAi cells (Green line). Failed constraint in wing discs (*br^disc^* ON) would also cause increased levels in RNAi cells (Magenta line). (C) Confocal images of positive (*E93-RNAi*) and negative (*lexA-RNAi*) controls. Yellow arrows indicate regions of RNAi expression. Stock identification numbers are indicated (see **Table S1**). (D) Quantification of changes in *br^disc^*reporter activity induced by RNAi. Boxplots summarize ratios of *br^disc^*signal in RNAi cells to WT cells. Each datapoint is a different wing. Each RNAi line tested is plotted on the y-axis, with gene symbol followed by RNAi line ID. “v” preceding line number indicates VDRC. A negative control *lexA* RNAi (magenta) has a ratio of ∼1. Subunits of the BAP complex are indicated in teal. Inset illustration depicts method of quantification. X-axis is log2 transformed. (E) Confocal images of *br^disc^*activity after RNAi of select core components of the BAP complex. Images are maximum projections. Scale bars are 100µm.

We focused our RNAi screen on nucleosome remodelers, reasoning that the functions of these enzymatic complexes in controlling nucleosome occupancy, positioning, and stability may contribute to developmentally programmed changes in enhancer accessibility. We tested a total of 49 RNAi lines corresponding to 31 genes encoding components of major ATP-dependent nucleosome remodeling complexes (Clapier *et al*. 2017; **Table S1**). These include members of all four families of remodeling complexes: Imitation Switch (ISWI), Switch/Sucrose Non-Fermenting (SWI/SNF), Chromodomain Helicase DNA-binding (CHD), and Inositol requiring 80 (INO80). Specific *Drosophila* nucleosome remodeling complexes include members of the ACF complex, the Brahma Complex (BAP and PBAP), the Chromatin Accessibility Complex (CAC), the Domino Complex, the INO80 complex, the Nucleosome Remodeling Deacetylase (NuRD) complex, the Nucleosome Remodeling Factor (NURF) complex, the Toutatis-containing chromatin Remodeling Complex (TORC), and several additional non-complex associated SNF2-like remodeler proteins (**Table S1**). To summarize the results of the screen we quantified the average intensity of the *br^disc^* reporter in RNAi expressing cells (anterior compartment), normalized to WT cells (posterior compartment) within the same wing (KD/WT) (**Fig 2D**). 2 genes were identified that decreased reporter activity in pupal wings, including Iswi, which is a component of the ACF, CHRAC, and NURF remodeling complexes (Bouazoune and Brehm 2006; **Fig S2C**). 8 genes were identified that increased reporter activity (**Fig 2D**; **Table S1**). Remarkably, 5 of these 8 genes are subunits of the *Drosophila* SWI/SNF BAP complex. These include, *osa*, *moira* (*mor*), *Snr1*, *Bap111*, and the core ATP-ase Brahma (*brm*). In some cases, we found that RNAi lines targeting the same gene gave divergent results in our screen. For instance, two independent RNAi lines for *Snr1* yielded some of the strongest increases in *br^disc^* activity (lines 32372 and 108599) while a third line (67929) produced little change. Similarly, of the two RNAi lines targeting *brm*, only one (31712) had a significant effect on *br^disc^* reporter activity. Notably, RNAi lines that significantly impacted reporter activity often caused lethality, with many animals dying as pupae or pharate adults (**Table S1**). By contrast, the *brm* and *Snr1* RNAi lines that did not impact reporter activity had little to no impact on animal survival or wing development. Since both *brm* and *Snr1* are essential genes, we interpret the lack of phenotype caused by these RNAi lines to be a consequence of poor target knockdown. Due to the enrichment of Brahma complex members among hits, we chose to characterize the role for this nucleosome remodeler in *br^disc^* repression. We did not pursue other hits further.

There are two distinct versions of the Brahma complex in *Drosophila*, BAP and PBAP, which are defined by the mutually exclusive association of either Osa (BAP), or Polybromo, SAYP, and Bap170 (PBAP, Cenik and Shilatifard 2021). We find that multiple RNAi lines targeting *osa* resulted in de-repression of *br^disc^* in the pupal wing, whereas three independent RNAi lines for *polybromo* had no effect on the normal dynamics of *br^disc^*by either qualitative observation or image quantification (**Fig 2E; Fig S2D**). We did not observe significant lethality or dramatic changes in wing morphology with any of the tested *polybromo* RNAi lines, suggesting that either these *polybromo* RNAi reagents are ineffective in the context of our screen or Polybromo is not required for wing development at this stage. Although we cannot definitively exclude PBAP, the finding that Osa is required for *br^disc^* reporter repression demonstrates a role for the Osa-specific BAP complex in the dynamic regulation of this enhancer. Homozygous *osa* mutant cells generated by mitotic recombination also exhibited increased *br^disc^* reporter activity (**Fig S2E**), further supporting a role of Osa and the BAP complex in *br^disc^* repression. Accordingly, we focused on the Osa-specific BAP complex in subsequent experiments.

### Osa is largely dispensable for pupal patterns of chromatin accessibility

Deactivation of temporally dynamic enhancers is associated with decreased chromatin accessibility over developmental time, and failure to deactivate temporally dynamic enhancers coincides with aberrantly persistent chromatin accessibility (Uyehara *et al*. 2017). Our observation that Osa depletion causes de-repression of the *br^disc^* reporter in pupal wings (**Fig 2E**) raised the possibility that the BAP complex may be required for closing of temporally dynamic enhancers. To test the genome-wide role of Osa in developmental control of chromatin accessibility, we performed FAIRE-seq in an *osa* degradation genotype. We employed the GFP deGrad system in conjunction with a genotype, *osa^GFP^*, in which both *osa* alleles are tagged with GFP (hereafter: Osa-deGrad; Buszcak *et al*. 2007, Caussinus *et al*. 2012; **Fig S3A**). The GFP deGrad system enables target proteins to be rapidly degraded, which is especially important in pupal wings because they undergo few cell divisions following pupariation (Ma *et al*. 2019, **Fig 3A**). Animals homozygous for *osa^GFP^* are viable and do not exhibit any morphological or developmental defects, indicating that Osa-GFP protein is functional. Confocal microscopy of *osa^GFP^* heterozygous imaginal wing discs showed nuclear-localized GFP that colocalized with endogenous Osa. Moreover, GFP signal was specifically depleted upon expression of Osa RNAi, validating the identity of this genotype (**Fig S3B**). Osa-GFP degradation was induced in animals homozygous for *osa^GFP^* during the late larval stage (3LW). Prepupal animals were staged (0-12h APF) and aged for ∼24h before dissection such that total degradation time was ∼36h (**Fig 3A**). Consistent with our RNAi results, we observed near-complete loss of Osa-GFP protein in the pupal wing blade under these conditions and a corresponding increase in *br^disc^*reporter activity, demonstrating the efficacy of Osa-deGrad depletion (**Fig 3B**). Immunofluorescence for Osa under these degradation conditions confirmed significantly reduced nuclear Osa signal relative to control (**Fig S3C**). Osa-deGrad flies that were permitted to develop longer exhibited reduced wing size and high rates of lethality, with most animals dying as pharate adults.

**Figure 3.**
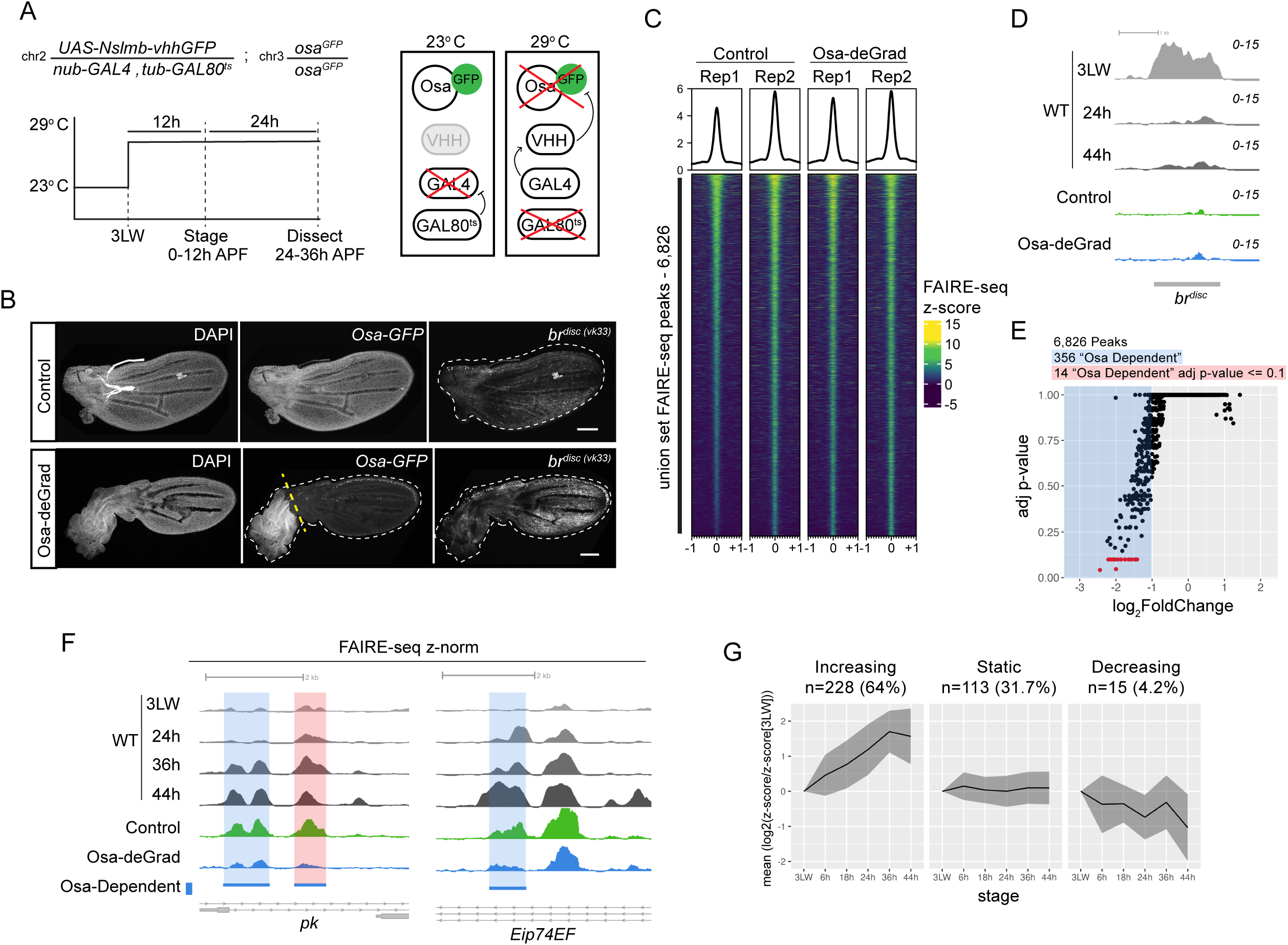
Osa is not required to close *br^disc^* and is dispensable for pupal chromatin accessibility patterns. (A) Illustrations of Osa^GFP^ degradation (Osa-deGrad) genotypes and experimental design. (B) Confocal images of Osa-deGrad experimental genotypes. Yellow dashed line indicates where wings were cut during sample collection. Scale bars are 100µm. Images are maximum intensity projections. (C) Heatmaps and average signal plots of z-normalized FAIRE signal within the union set of FAIRE peaks from Control and Osa-deGrad pupal wings. Plotted range is +/-1kb from peak center. Peaks are ranked by signal in Control Rep1. (D) Browser shot of z-normalized FAIRE signal from Osa-deGrad (blue), Osa-deGrad Control (green), and WT (grey) imaginal wing discs at the endogenous *br^disc^* enhancer. (E) Scatterplot of log_2_FoldChange of Osa-deGrad/Control FAIRE-seq signal (x axis) relative to adjusted p-value (y axis). Peaks with log_2_FoldChange <= –1 (Osa-dependent) are highlighted in blue. Peaks with an adjusted p-value <= 0.1 are colored red. (F) Browser shot of FAIRE signal at representative “Osa-dependent” sites (blue bars and highlights, red highlight indicates adjusted p-value < 0.1) near the *prickle* (*pk*) and *Eip74EF* loci. (G) Line plots of the average WT FAIRE log_2_FoldChange relative to 3LW, with standard deviation as grey ribbon, within 356 “Osa-dependent” sites. Sites are split by whether they increase in accessibility relative to the 3LW stage (“Increasing”), have little change (“Static”), or lose accessibility (“Decreasing”) (see Methods). The x axis denotes stages of wing development from 3LW to 44h APF. All z-normalized FAIRE signal in browser shots are pooled replicates.

Despite the strong impact of Osa-deGrad on wing development, *br^disc^*reporter activity, and survival, FAIRE-seq revealed minimal changes in chromatin accessibility profiles relative to controls samples, including at *br^disc^*(**Fig 3C,D**). A union set of 6,826 open chromatin peaks was identified between both control and Osa-deGrad samples (**Fig S3D**). Pearson correlation coefficients of z-normalized FAIRE signal revealed high correlation between both replicates of control and Osa-deGrad pupal wing profiles, indicating that Osa degradation minimally affects open chromatin profiles (**Fig S3E**). We conclude that increased *br^disc^* activity observed in *osa* loss-of-function pupal wings is not due to failure to close the *br^disc^* enhancer. These findings also demonstrate that developmentally programmed opening and closing of wing enhancers occurs normally genome wide in the absence of Osa.

Previous studies in mammalian experimental systems have observed a role of Brahma Complex orthologs in promoting chromatin accessibility (Kelso *et al*. 2017, Hendy *et al*. 2022). To test for the possibility of subtle changes in accessibility, we compared FAIRE signal between Osa-deGrad and control samples and observed a unidirectional skew toward lower FAIRE signal in Osa-deGrad relative to control (**Fig 3E**). We note that while we find 356 regions (5.2%) with reduced accessibility (log_2_FoldChange <= –1, “Osa-dependent”) following Osa-GFP degradation, only 14 (0.2%) were found to be statistically significant (adj p-value <= 0.1) due to variability between replicates, raising the possibility that some of these regions are false positives. Examples of these Osa-dependent sites occurred at genes including *prickle* (*pk*) and the ecdysone response gene *Eip74EF* (**Fig 3F**). Both of these Osa-dependent sites exhibit temporally dynamic accessibility, with low accessibility observed in larval wing imaginal discs becoming progressively more open later in pupal stages. To determine if temporally dynamic accessibility is a general feature of Osa-dependent FAIRE peaks, we categorized each peak as either ‘Increased’, ‘Static’, or ‘Decreased’ based on the wild-type FAIRE-seq signal at that site during pupal stages relative to the late larval stage (see **Methods**). We find that 64% (228/356) of Osa-dependent sites correspond to regions that increase in accessibility between larval and pupal stages (**Fig 3G**), whereas only 4.2% of Osa-dependent sites decrease in accessibility over the same time interval. This finding suggests that while the effect of Osa-GFP degradation is minor, the subtle losses in accessibility observed are most often associated with regions that open between early and late wing development. Collectively, we conclude that Osa is not required for large, binary changes in “open” or “closed” chromatin over time during wing development. Instead, it is required for only a small number of sites to achieve full accessibility.

### The *br^disc^*reporter is active in a small number of pupal wing cells upon Osa loss of function

Our FAIRE-seq data indicate that Osa is not required for enhancer closing between early and late stages of wing development. The finding that *br^disc^* is closed in *osa* loss-of-function pupal wings raised the possibility that the enhancer is inactive despite the apparent increase in reporter activity. To directly test whether the *br^disc^* enhancer is active in *osa* loss-of-function pupal wings, we utilized the dual-fluorophore *br^disc^-switch* reporter (**Fig 1A**). We first depleted Osa using the same RNAi-mediated knockdown conditions employed in the nucleosome remodeler screen, but using the posterior *en-GAL4, tub-GAL80^ts^* (*en^ts^*) driver. The switch from tdTomato to myr-GFP reporter output was induced at a timepoint after *br^disc^* deactivation (>24h APF). Under these conditions, we found that the enhancer remains inactive in the great majority of pupal wing cells, with a few notable exceptions. Whereas control lexA knockdown pupal wings exhibited nascent GFP along the margin and in the posterior crossvein, similar to the enhancer’s pattern of activity in wild-type pupal wings (**Fig 1B**), *osa* knockdown pupal wings also exhibited nascent GFP expression in a subset of cells in the wing blade (**Fig 4A**). Notably, the membrane localized myr-GFP of the *br^disc^-switch* reporter revealed that wing blade cells in which *br^disc^* was active exhibited a distinct morphology resembling shaft cells of adult sensory organs. These were similar in appearance to the shaft cells located along the wing margin in which the *br^disc^* reporter normally reactivates during later pupal stages in wild-type animals (**Fig 4A Inset**). Elevated levels of nascent GFP were also observed in the posterior margin of *osa* knockdown pupal wings (**Fig 4A**). The detection of nascent GFP after *br^disc^* normally closes and deactivates indicates that *osa* knockdown causes the enhancer to be inappropriately active in a small number of cells. However, *br^disc^* activity was not detected in most cells that exhibited increased reporter levels in the initial nucleosome remodeler screen. It is possible that the small number of additional cells in which *br^disc^* is active in *osa* knockdown pupal wings is too few to impact whole wing open chromatin profiles, thus explaining the closed appearance of the enhancer in our FAIRE-seq data.

**Figure 4.**
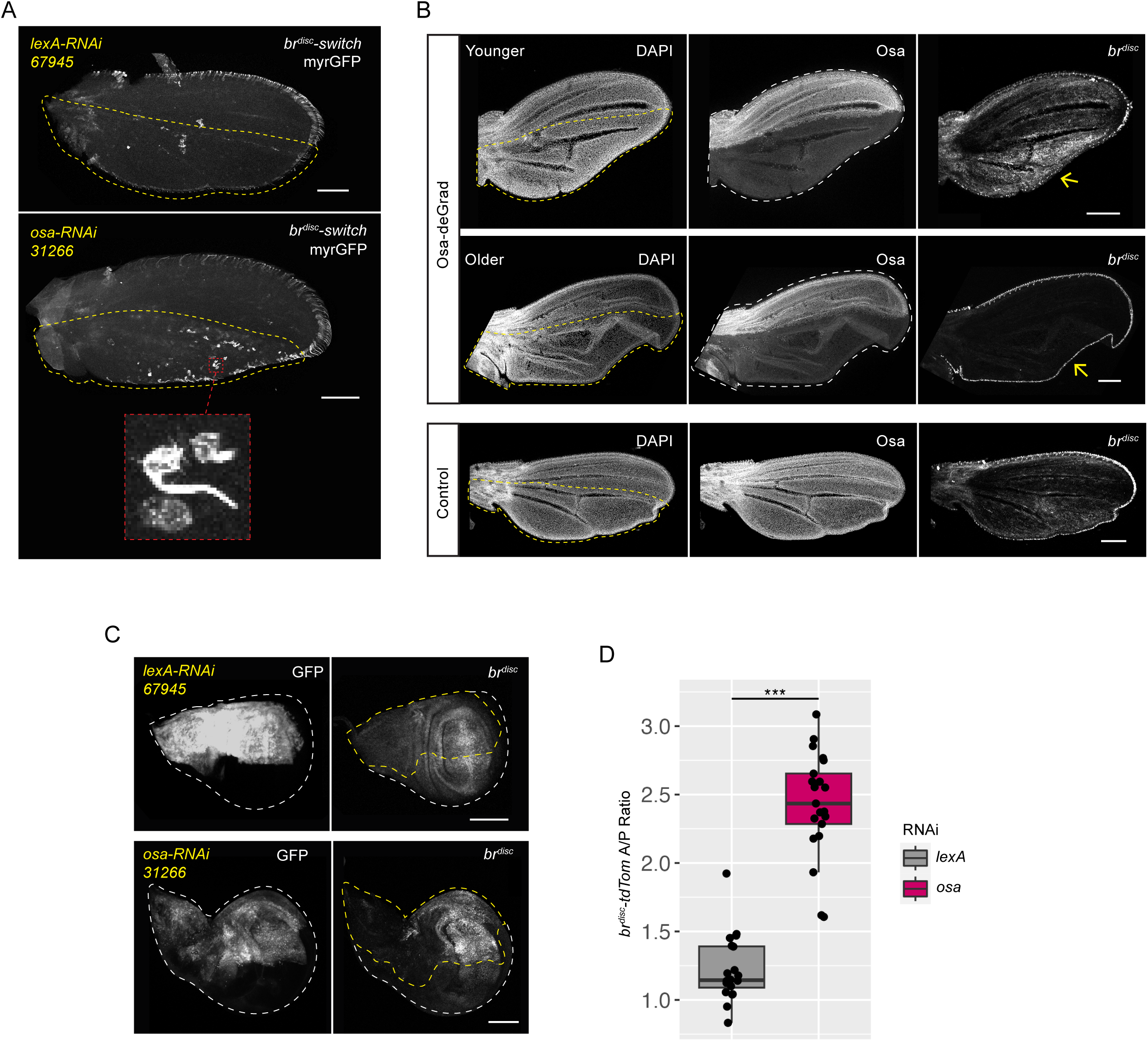
Osa is required to constrain *br^disc^* activity in wing imaginal discs. (A) Confocal images of *br^disc^-switch* nascent myr-GFP signal in the pupal wing in negative control *lexA* RNAi or *osa* RNAi. (B) Confocal images of *br^disc^-tdTomato-PEST* activity in 30-42h APF wings following late induction of Osa-deGrad. Approximate regions of Osa degradation are outlined with yellow dashed line in the DAPI channel. “Younger” indicates a wing closer to 30h APF of age. “Older” indicates a wing closer to 42h APF (see Methods). A negative control in which Osa-deGrad was induced in an *osa^GFP^*heterozygote (*osa^GFP^/osa*) is shown for comparison (Control). Yellow arrows denote regions of differential reporter activity for comparison. (C) Confocal images of *br^disc^* activity in *osa-RNAi* and control *lexA-RNAi* wing imaginal discs. GFP marks domain of RNAi expression (outlined by dashed yellow line). (D) Quantification of *br^disc^* reporter increase in response to *osa*-*RNAi* in wing imaginal discs, compared to control *lexA*-*RNAi*. The y-axis is a ratio of *br^disc^* signal in the anterior (RNAi-expressing) versus the posterior (WT) cells. Asterisks indicate significance (*** = p-value < 1e-13, Two-sample *t*-test). Images are maximum intensity projections. Scale bars are 100µm.

Sensory organs do not normally develop within the wing blade. In wild-type tissues, sensory organ precursors (SOPs) are specified with stereotypical spatial and temporal patterns, with the last SOPs in the wing being specified during prepupal stages. Once specified, SOPs undergo two rounds of cell division and fate specification, resulting in development of a single shaft, socket, sheath, and neural cell, which together compose an adult sensory organ (Couso *et al*. 1994; Furman and Bukharina 2012). The appearance of *br^disc^* activity in cells with shaft-like morphology in the wing blade indicated that *osa* knockdown leads to development of ectopic sensory organs. Consistent with this hypothesis, *osa* knockdown also resulted in ectopic expression in the wing blade of Elav, a marker of neural cell identity (**Fig S6**). This finding is in agreement with prior studies in which combinations of *osa* hypomorphic alleles and loss-of-function clones caused ectopic sensory organ development (Heitzler *et al*. 2003; Terriente-Félix and de Celis 2009). We speculated that by initiating RNAi expression in larval wing discs, early loss of *osa* function leads to ectopic sensory organ development accompanied by *br^disc^* activation. To test this hypothesis, we sought to knockdown *osa* function later in wing development, reasoning that the delay would reduce the likelihood of disrupting development of sensory organs, which are determined by the end of prepupal stages (Couso *et al*. 1994). We returned to the Osa-deGrad system due to its rapid depletion of Osa protein, in combination with the *en^ts^* driver. Osa degradation was initiated in 0-12h prepupae, ∼48h later than initiation of knockdown in the RNAi screen. Examination of *br^disc^* reporter activity 30h later revealed two phenotypic classes that correlated with wing age. In younger pupal wings, there was clear de-repression of *br^disc^*in Osa-deGrad cells relative to wild-type cells in the same wing. By contrast, older pupal wings exhibited no sign of *br^disc^* de-repression in the wing blade (**Fig 4B**). We interpret these findings as being a consequence of the developmental stage when Osa degradation was initiated. Since the duration of Osa depletion was the same for both phenotypic classes, the younger pupal wings, which exhibit *br^disc^* de-repression, would have been at an earlier developmental stage when Osa depletion initiated than the older pupal wings, which do not exhibit *br^disc^* de-repression. We conclude that Osa is not required for *br^disc^* deactivation. Instead, any detected increase in reporter activity is likely due to indirect consequences stemming from disruption of *osa* function early in sensory organ development.

### Osa is required to constrain *br^disc^* reporter activity in wing imaginal discs

We hypothesized that if Osa and other BAP complex members are required for reduced *br^disc^* reporter levels in pupal wings, as indicated by our RNAi screen results, but they are not required for chromatin closing or for *br^disc^* reporter deactivation, then Osa may be required to repress *br^disc^* reporter activity at an earlier stage of development **(Fig 2B**). To test this hypothesis, we assayed *br^disc^* reporter activity in wing imaginal discs following Osa knockdown, which corresponds to a developmental stage when *br^disc^*is normally active. At this timepoint, wing discs are approximately two days younger than the pupal wings assayed in our RNAi screen. We observed a marked increase in reporter activity in *osa* knockdown cells relative to control cells. By contrast, a negative control RNAi targeting lexA did not affect *br^disc^* reporter activity at this stage. (**Fig 4C**). Quantification of the ratio of reporter signal in RNAi expressing versus control cells confirmed significant hyperactivation of the reporter in *osa* knockdown but not in control lexA knockdown cells (p-value=6.84e-14, Two-sample *t*-test; **Fig 4D**). The requirement of the BAP complex to constrain *br^disc^* activity in the wing disc was further validated by independent knockdown of a different BAP complex member *brm*, as well as an independent *osa* RNAi line (330350), both of which also resulted in increased *br^disc^* reporter activity (**Fig S4A,B**). Thus, loss of Osa function results in hyperactivation of the *br^disc^* enhancer in cells in which it is already active. Increased *br^disc^* reporter levels following knockdown of BAP complex members indicates that BAP is required to constrain activity of the enhancer in wing imaginal discs. Because we find no evidence that Osa is required to close the *br^disc^*enhancer in pupal wings, we interpret the increased reporter activity observed in pupal wings to be a consequence of persistent reporter fluorophore following enhancer hyperactivation in wing imaginal discs (**Fig 2B, magenta line**).

### Osa directly binds *br^disc^* in larval wing imaginal discs as well as thousands of putative enhancers genome wide

Hyperactivation of the *br^disc^* reporter in wing imaginal discs following degradation of BAP complex members could be due to a direct loss of BAP function at the enhancer, or an indirect consequence of dysregulation of other *br^disc^* inputs. To determine if the BAP complex directly binds *br^disc^*, we performed CUT&RUN for Osa using the *osa^GFP^* allele. We performed anti-GFP CUT&RUN in homozygous *osa^GFP^* and WT control female wing imaginal discs. We identified 2,150 Osa-GFP peaks (see **Methods**), the great majority of which (1,953) did not overlap control peaks (**Fig 5A,B**). We focused on this set of Osa-GFP-specific peaks (“Osa peaks”) for all subsequent analysis.

**Figure 5.**
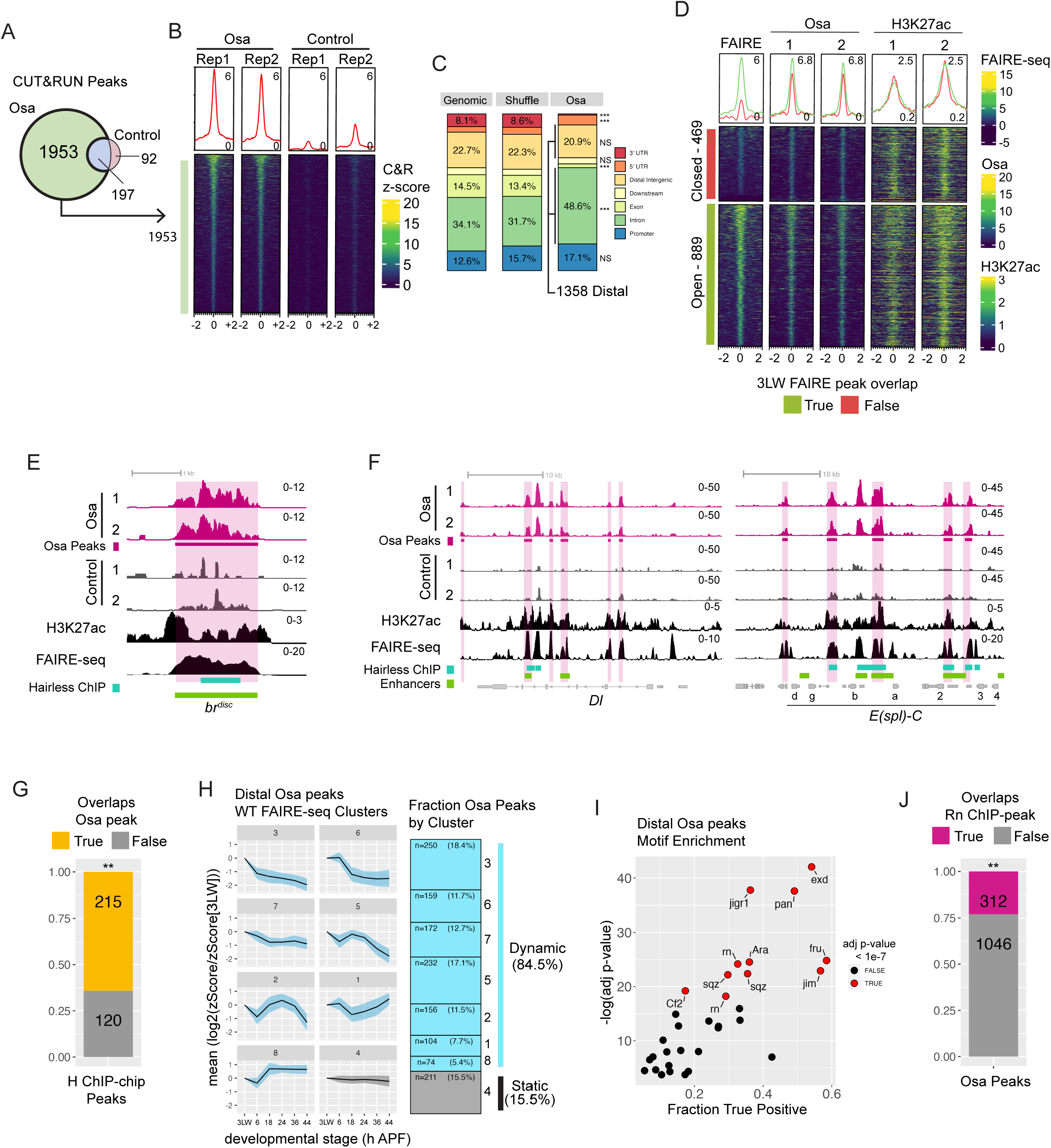
Osa directly binds *br^disc^* and thousands of putative enhancers in wing imaginal discs. (A) Venn diagram of peaks called in Osa-GFP (Osa) versus Control wing imaginal disc CUT&RUN experiments. (B) Heatmap and average signal plots of z-normalized CUT&RUN signal between experimental replicates within Osa-specific peaks. (C) Stacked barplots of the distribution of Osa peak genomic annotations relative to a 500bp-tiled genome-wide annotation (Genome), and a bootstrapped shuffle of Osa peaks (Shuffle). Asterisks indicate significance (*** = p-value < 0.0001, two-proportion z-test). (D) Heatmap and average signal plots of wing imaginal disc z-normalized FAIRE-seq, Osa CUT&RUN, and H3K27ac CUT&RUN signal within distal Osa peaks. Heatmaps are grouped by whether Osa peaks overlap a FAIRE peak in 3LW wing discs. (E,F) Browser shots of Osa CUT&RUN signal (magenta) versus control (grey), H3K27ac z-normalized signal (black), and WT FAIRE-seq (black). Coordinates for Osa peaks (magenta), Hairless ChIP peaks (teal), and annotated enhancers (green) are indicated. Browsers depict the *br^disc^* enhancer (E), and the *Dl* and *E(spl)-C* loci (F). (G) Bar plot showing fraction of Hairless ChIP peaks that overlap Osa peaks (not restricted to distal only). Asterisks indicate significance (** = adj p-value < 0.001, Fisher’s exact test). (H) Line plots of the ratio of wild-type wing FAIRE-seq signal in distal Osa peaks for each of six developmental stages relative to 3LW (log_2_). Osa peaks were placed into eight categories by k-means clustering of the wild-type FAIRE time course data. Standard deviation shown by blue ribbon. Stacked barplot depicts fraction of distal Osa peaks associated with each cluster. Dynamic clusters (1,2,3,5,6,7,8) are colored blue. Static cluster (4) is colored grey. (I) Scatterplot of motifs enriched in distal Osa peaks, plotted by –log(adj p-value) and fraction of true positive. Motifs with an adjusted p-value < 1e-7 are colored in red. (J) Bar plot of the fraction of distal Osa peaks that overlap Rotund (Rn) ChIP-seq peaks. Asterisks indicate significance (** = adj p-value < 0.001, Fisher’s exact test).

Genomic feature annotation revealed that a majority of Osa peaks were enriched in distal intergenic regions and introns relative to a shuffled Osa peak control annotation (**Fig 5C**). We found that Osa peaks were significantly more abundant in “Introns” (48.6%, p-value=6.4e-58), and “5’ UTRs” (6.3%, p-value=7e-5). Osa peaks were significantly less abundant in “Exons” (2.4%, p-value=5.7e-46) and “3’ UTRs” (0.8%, p-value=5e-35, two-proportion z-test). Collectively, these findings indicate that Osa is predominantly bound to non-coding regions of the genome (86.6% Promoter | Distal Intergenic | Intron), consistent with an expected role in gene regulation. In order to focus our analysis on *cis*-regulatory elements with potential roles as developmentally dynamic enhancers, we selected Osa peaks that lie distal to promoters for use in subsequent analysis (1,358 peaks, “distal Osa peaks”).

To evaluate the relationship between Osa binding and potential enhancer activity, we next examined the overlap between distal Osa peaks and open chromatin sites in wing imaginal discs (Uyehara *et al*. 2017). We found that most distal Osa peaks (65%, 889) were associated with a high-degree of chromatin accessibility, whereas 35% (469) of distal Osa peaks did not overlap a FAIRE peak in wing imaginal discs (**Fig 5D**). Notably, 25% (119) of these Osa-bound “closed” sites were identified as a FAIRE peak in at least one later stage of wing development (**Fig S5F**). Thus, 74.2% (1008) of distal Osa peaks are bound at regions that are either open in wing imaginal discs or will open subsequently during a later stage of wing development. To further examine the regulatory potential of distal Osa peaks, we performed CUT&RUN for histone H3 lysine 27 acetylation (H3K27ac), an epigenetic mark associated with active enhancers, in wing imaginal discs. We found an enrichment of H3K27ac signal at highly accessible distal Osa peaks (**Fig 5D**). Interestingly, we also found H3K27ac signal at distal Osa peaks that do not overlap a FAIRE peak, suggesting that some of these sites possess enhancer activity despite exhibiting low chromatin accessibility (**Fig 5D**). Together, the correlation between chromatin accessibility and H3K27ac enrichment at distal Osa peaks indicates these sites are likely to function as enhancers in developing wings.

To further define the relationship between Osa occupancy and regulatory DNA, we examined binding at previously characterized enhancers. Firstly, we found that Osa is bound at the endogenous *br^disc^* enhancer with broad signal observed across the entire enhancer in both replicates. By contrast, CUT&RUN signal apparent at the *br^disc^* enhancer in control experiments was non-reproducible and restricted to narrow regions, which we interpret as being due to opportunistic MNase digestion of this highly accessible DNA (**Fig 5E**). The presence of Osa at the endogenous *br^disc^* enhancer strongly suggests that direct binding of the BAP complex to *br^disc^* is required to constrain its activity.

In addition to *br^disc^*, we observed Osa bound at multiple genes known to be regulated by the Brahma Complex during wing development, such as at components of the *Drosophila* Notch signaling pathway. There is a well-established connection between Brahma complexes and Notch signaling. Mutants of Notch pathway genes enhance *brm* dominant negative allele phenotypes, Osa loss of function increases expression of the proneural Notch targets *achaete* and *scute* (*ac/sc*), and both Brm and Mor are required for full induction of the Notch target genes in the *Enhancer of Split complex* (*E(spl)-C*) locus (Elfring *et al*. 1998; Heitzler *et al*. 2003; Armstrong *et al*. 2005; Pillidge and Bray 2019). Consistent with this relationship, we observed high-amplitude Osa binding sites at the genes encoding the Notch ligands *Delta* (*Dl*) and *Serrate* (ser), the Notch receptor *Notch* (*N*), and Notch target *E(spl)-C* genes (**Fig 5F, S5A-C**). At least two of the Osa peaks in the *Dl* locus corresponded to previously characterized enhancers, including the *Dl^SOP^* enhancer, which is active within sensory organ precursor cells in wing imaginal discs, and the *Dl^teg^*enhancer, which is active in the tegula, hinge, and anterior notum (Uyehara and McKay 2019). Osa binding in the *E(spl)-C* locus overlaps the mα, mβ, m2, and m3 enhancers, which contribute to proneural cluster development in wing imaginal discs. Like many signaling pathways, Notch signaling relies on the action of co-repressors to limit expression of Notch targets in the absence of signal. In *Drosophila,* the co-repressor Hairless binds the Notch signaling effector Suppressor of Hairless (Su(H)) and has been found to bind hundreds of sites across the genome in the wing disc, including known regulatory sites that require Hairless for negative regulation (Chan *et al*. 2017). Using previously published Hairless ChIP data from wing imaginal discs, we examined the correlation between Osa and Hairless binding (Chan *et al*. 2017). We found that the majority (64.2%) of Hairless peaks intersect an Osa binding site, indicating a significant overlap between these gene regulatory proteins (adj p-value=9.99e-4, Fisher’s exact test; **Fig 5G**). Notably, we also observed that Hairless was bound at the endogenous *br^disc^* enhancer, as well as at known and putative enhancers of the *Dl* and *E(spl)-C* loci (**Fig 5E,F**). These findings further support the strong association between the BAP complex and Notch signaling. More generally, the concerted presence of Osa binding at *bona fide* enhancers indicates that the BAP complex is a direct regulator of transcriptional programs with major roles in wing development.

In addition to binding known regulatory elements, we found that Osa binding is also correlated with genomic loci that exhibit temporal changes in chromatin accessibility. A previous FAIRE-seq time course of wild-type wings identified distinct patterns of temporal accessibility changes (Uyehara *et al*. 2017; Nystrom *et al*. 2020; **Fig 5H**, **S5E**). By clustering FAIRE signal in Osa-bound regulatory sites, we found that 84.5% of Osa-bound distal regulatory sites were associated with FAIRE-seq peaks that exhibited temporal changes during wing development, whereas only 15.5% of peaks exhibited static accessibility (**Fig 5H**). The correlation between distal Osa peaks and dynamic rather than static accessibility, indicates that Osa is associated with regulatory regions that are likely stage-specific and are either being actively used at larval stages or possibly constrained from being used until a later stage.

Like most nucleosome remodeling complexes, the BAP Complex does not exhibit sequence-specific DNA binding but is instead thought to be recruited to target loci by transcription factors. Osa contains an AT-Rich Interacting Domain (ARID) that facilitates interaction with DNA, but this domain has been shown to confer little to no sequence preference on BAP complex DNA binding (Collins *et al*. 1999; Patsialou *et al*. 2005). To identify candidate factors that contribute to the recruitment of Osa to its binding sites, we performed motif enrichment analysis of sequences around distal Osa peaks. Of the highest significance motifs, several were associated with major signaling pathways and wing patterning programs. Notably, we found motifs for the homeodomain factors Extradenticle (Exd) and Araucan (Ara), the Wingless-signaling effector Pangolin (Pan), and the zinc-finger transcription factors Squeeze (Sqz) and Rotund (Rn) (**Fig 5I**). Enrichment of Pan motifs in Osa binding sites is notable because Osa has been proposed to repress Wingless target genes (Collins and Treisman 2000). Our findings indicate that this repression could be direct. For instance, the Wingless target gene, *nubbin*, which is ectopically expressed in *osa* mutants, has several Osa binding sites in wing imaginal discs (Collins and Treisman 2000; **Fig S5D**). The enrichment of Pan motifs and others in Osa binding sites suggests that the BAP complex is broadly utilized by the major signaling pathways and patterning factors that shape wing development. To extend this observation further, we examined recently published Rn ChIP-seq data from wing imaginal discs and found that 23% (312) of distal Osa peaks overlap with a Rn peak, which is greater than expected by chance as tested by overlap with shuffled Osa peak controls (adj p-value=9.99e-4, Two-sample *t-*test) (**Fig 5J**). This correlation between Rn and Osa binding at regulatory sites in the wing disc further supports the connection between Osa and active wing developmental programs.

### Osa is required to constrain Delta activation in wing imaginal discs

Notch signaling performs multiple critical roles in wing imaginal discs. Notch-mediated lateral inhibition is necessary for selecting the sensory organ precursor cells that form the chemosensory and mechanosensory organs of the adult wing. Notch signaling also initiates specification of wing vein cell fates. Both of these processes depend on patterned expression of the Notch ligand, *Dl*, which has an extensive *cis*-regulatory domain. It has been previously observed that Osa is involved in the regulation of *Dl* expression in parts of the wing disc pouch (Terriente-Félix and de Celis 2009). Due to the discovery of multiple Osa binding sites at known and putative enhancers within the *Dl* locus, we hypothesized that Osa regulates *Dl* expression directly. Dl is expressed in larval wing discs in two rows of cells flanking the dorsal/ventral (DV) boundary of the developing wing margin, and in perpendicular stripes marking the developing veins (Doherty *et al*. 1996). We found that depletion of Osa from the anterior compartment of the wing disc resulted in increased Dl levels and subtle expansion of the Dl pattern most notably around the L2 provein stripe (**Fig 6A**). To support this observation, we quantified Dl levels around the margin in both Osa and lexA control knockdown experiments. For each wing disc, measurements around the RNAi expressing anterior margin were normalized to non-RNAi expressing cells in the posterior margin (see Methods). We find that Dl levels were significantly higher in Osa knockdown relative to control (p-value=9.67e-11, Two-sample *t*-test, **Fig 6B**). Notably, our observation that Osa negatively regulates Dl expression is in disagreement with previous work that found Osa depletion leads to reduced Dl in the L3 and L4 proveins (Terriente-Félix and de Celis 2009; see **Discussion**). Together, these findings demonstrate that enhancer hyperactivation in the absence of the BAP complex is correlated with increased expression of the Notch ligand.

**Figure 6.**
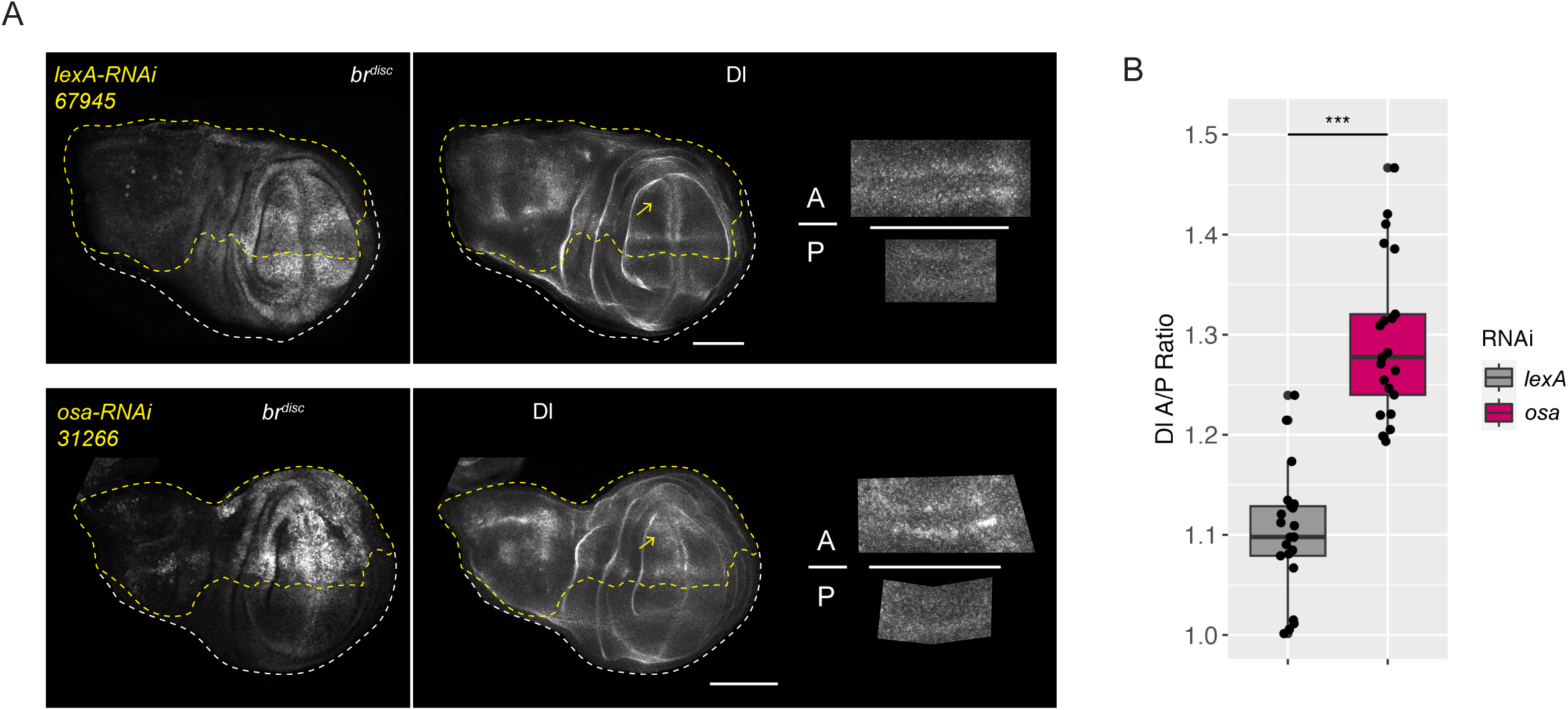
Osa negatively regulates Delta expression. (A) Confocal images of *br^disc^* reporter activity and immunofluorescence of Delta protein in wing imaginal discs from negative control *lexA* or *osa* RNAi. RNAi expressing cells are outlined with yellow dashed line. Insets show representative ROI selections in the anterior (“A”) and posterior (“P”) used for quantification (see **Methods**). Yellow arrows highlight expansion of Dl pattern around L2 provein relative to control. (B) Image quantification of Delta levels in lexA control and Osa RNAi experiments. Asterisks indicate significance (*** = p-value < 1e-10, Two-sample *t*-test). Maximum intensity projections are shown. Scale bars are 100µm.

## DISCUSSION

We set out to investigate possible roles of nucleosome remodeling complexes in developmentally programmed enhancer regulation, with a particular focus on enhancer closing and deactivation. Using reporters of the previously characterized and developmentally dynamic wing enhancer, *br^disc^*, we performed an *in vivo* RNAi screen that identified members of the *Drosophila* SWI/SNF (BAP) complex as repressors of enhancer reporter activity. Surprisingly, we find that the BAP-specific subunit Osa is not required to close *br^disc^* and is globally dispensable for binary changes in accessibility, closing or opening, between early and late stages of wing development (**Fig 3**). Rather than being required for enhancer deactivation, we instead find that Osa is required to constrain activity of the *br^disc^* enhancer when it is in the ON state in wing discs (**Fig 4**). Genome-wide profiling of Osa binding revealed that Osa bound extensively to sites with signatures of active regulatory DNA (open and H3K27ac enriched), including at multiple known and putative enhancers of Notch pathway component genes (**Fig 5**). Analysis of binding sites of Osa and the Notch co-repressor Hairless revealed significant co-enrichment of these proteins genome wide, suggesting a direct coregulatory relationship between Notch signaling responses and the BAP complex. Finally, we find that Osa depletion in wing discs leads to upregulation of the Notch ligand, Delta, further supporting a central role of Osa and the BAP complex in regulating Notch pathway activity (**Fig 6**).

### Is the BAP complex required for control of chromatin accessibility in developing *Drosophila* wings?

Thousands of *cis*-regulatory elements exhibit chromatin accessibility changes during the first two days of pupal wing development in *Drosophila*, which drive the dynamic gene expression changes that underlie progressive determination of cell fates (Uyehara *et al*. 2017; Ma *et al*. 2019). We hypothesized that nucleosome remodeling complexes work with sequence-specific transcription factors to bring about these kilobase-sized transitions in chromatin state. However, we observed no requirement for Osa in either opening or closing enhancers genome-wide (**Fig 3**). This is a surprising finding because SWI/SNF complex function has been found to be required for proper control of chromatin accessibility in mammalian cells. For example, genetic removal or chemical inhibition of the SWI/SNF ATPase Brg1 in mouse embryonic stem cells results in loss of accessibility genome wide (Iurlaro *et al*. 2020). Similarly, loss of the Osa ortholog ARID1A, which is commonly mutated in cancers (Kadoch *et al*. 2013), results in both loss and gain of open chromatin sites in human cells (Kelso *et al*. 2017). Whereas we observed subtle decreases in accessibility at a subset of open chromatin sites upon Osa knockdown, we did not find evidence for a global role of Osa in binary chromatin state transitions from closed to open or open to closed, leading us to conclude that Osa is not required for these developmentally programmed epigenetic changes. An alternative explanation is that our methods did not sufficiently deplete Osa below a minimal threshold. We disfavor this possibility because no Osa-GFP signal remains after nanobody-mediated degradation, and immunostaining with Osa antibodies likewise revealed little nuclear signal above background (**Fig S3**). Moreover, we observed developmental phenotypes consistent with Osa loss of function. Another possible explanation is that the role of the BAP complex in regulating chromatin accessibility is compensated for by the PBAP complex. Synthetic lethal phenotypes caused by perturbation of subunits from distinct SWI/SNF complex subtypes have been reported, supporting the potential of functional redundancy (Michel *et al*. 2017; Helming *et al*. 2014; Wilson *et al*. 2015). Lastly, multiple nucleosome remodelers can be found at the same genomic targets (Morris *et al*. 2014), raising the possibility of compensation by other complexes.

### What is enhancer constraint?

SWI/SNF nucleosome remodeling complexes were first identified for their role in counteracting Polycomb-mediated repression and establishing regions of nucleosome depletion in order to facilitate transcription (Kassis *et al*. 2017; Cenik and Shilatifard 2021). Subsequent work has demonstrated that SWI/SNF complexes are required to maintain nucleosome depleted regions, high levels of H3K27ac, and enrichment of the histone variant H3.3 at enhancers and promoters (Alver *et al*. 2017; Schick *et al*. 2021; Blumli *et al*. 2021; Weber *et al*. 2021; Hendy *et al*. 2022; Reske *et al*. 2022). In addition to their role in gene activation, SWI/SNF complexes have also been implicated in gene repression (Treisman *et al*. 1997; Moshkin *et al*. 2007; Zraly *et al*. 2012; Kelso *et al*. 2017; Weber *et al*. 2021), including repression of Wingless target genes during wing development (Collins *et al*. 2000). Here, we find that the BAP complex constrains activity of the developmentally dynamic *br^disc^* enhancer, but it is not required for closing or deactivation. DNA binding profiles reveal that Osa binds the *br^disc^*enhancer while it is active in developing imaginal wing discs, suggesting its role in enhancer constraint is direct. How might SWI/SNF function to achieve constraint? SWI/SNF complexes are generally understood to slide and/or eject nucleosomes by translocating DNA around the histone octamer (Clapier *et al*. 2017). Nucleosome mobilization could result in repression if DNA translocation blocked a binding site for an activator. Conversely, increased accessibility mediated by SWI/SNF could uncover a repressor binding site. Differential accessibility of repressor binding sites in a wing spot enhancer was recently proposed as a mechanism involved in morphological diversification between *Drosophila* species (Ling *et al*. 2023). Another possible direct mechanism is through changes in histone acetylation via collaboration with the NuRD complex. A recent study in human endometriotic epithelial cells demonstrated that the Osa ortholog ARID1A is required to maintain levels of the histone variant H3.3 at active enhancers, which in turn is required to recruit NuRD complex components and limit the accumulation of active H3K27ac levels (Reske *et al*. 2022). Lastly, iterative cycles of nucleosome remodeling activity driven by ATP hydrolysis could impact the dynamics of transcription factor occupancy at target enhancers, which in turn could impact their potency as transcriptional regulators (Morris *et al*. 2014; Brahma and Henikoff 2023). In addition to these direct mechanisms, it is also possible, though not mutually exclusive, that SWI/SNF-dependent enhancer constraint is an indirect consequence of SWI/SNF-dependent repressor activation. For example, failure to activate the transcriptional repressors encoded by the *Enhancer of split complex* locus could contribute to hyperactivation of Notch pathway target genes in *Osa* loss of function wings (see below).

### The BAP complex as a direct regulator of Notch signaling

Our findings point to an important role of Osa in Notch pathway function. This is supported by prior studies that have discovered strong regulatory connections between the Notch pathway and the BAP complex. Genetic screens found that alleles of *Dl* dominantly enhance phenotypes of an ATP-ase dead *brm* allele (*brm^K804R^*; Armstrong *et al*. 2005). BAP complex members have also been found to regulate the expression of Notch signaling targets, such as genes encoded by the *Enhancer of split complex* and *achaete/scute* loci (Armstrong *et al*. 2005; Pillidge *et al*. 2019). Our genomic profiling of Osa in wing imaginal discs revealed clusters of Osa binding at putative regulatory sites at loci encoding the Notch ligands Dl and Ser, at the gene encoding the *Notch* receptor itself, and at enhancers of the *Enhancer of split complex* (**Fig 5, S5**). Interestingly, it has been previously reported that Osa negatively regulates expression of the proneural genes *achaete* and *scute*, but we observed little binding of Osa around these genes sparing a single potential binding site that also has a relatively high degree of signal in negative controls (Armstrong *et al*. 2005). This suggests the regulation of *achaete* and *scute* by the BAP complex may be indirect. In addition to extensive binding of Osa at genes encoding Notch pathway components, we also find significant co-enrichment of Osa binding and the Notch pathway co-repressor Hairless, including at the *br^disc^* enhancer **(Fig 5G**). Thus, the BAP complex may directly regulate Notch target genes genome wide. Together, our binding data strengthen the previously observed regulatory relationship between the BAP complex and Notch signaling.

Several observations made through the course of our study suggest a regulatory connection between the *br^disc^* enhancer, the BAP complex, and the Notch signaling pathway. The *br^disc^* enhancer may itself be a Notch pathway target gene. In addition to being bound Hairless, the pattern of *br^disc^* activity in wing imaginal discs suggests positive input from Notch signaling. The highest levels of enhancer activity in the pouch of wing imaginal discs are typically observed along the presumptive wing margin and in two dorsal-ventral stripes that extend away from the margin that resemble the wing proveins (**Fig 4C**). Each of these regions overlap high levels of Dl expression. The activity of *br^disc^* in pupal wings is also suggestive of Notch pathway input. *Br^disc^* is reactivated in the sensory organs located along the wing margin approximately 40hAPF. Notch signaling is required for determining the fates of these sensory organ cells. Moreover, sensory organ development is particularly sensitive to the levels of Notch pathway signaling, with too much or too little Notch signaling leading to sensory organ developmental defects. Hyperactivation of the Notch pathway may also explain development of ectopic sensory organs and activation of the *br^disc^*enhancer in shaft cells of the developing pupal wing blade upon Osa loss of function. Collectively, these observations suggest that the *br^disc^* enhancer is responsive to Notch signaling, and that the BAP complex may be required to directly constrain Notch target gene activity, possibly in collaboration with Hairless. A lack of proper constraint by the BAP complex at enhancers of Notch signaling component genes and of Notch target genes may result in the observed development of ectopic of bristles and neurons (**Fig S6**). This possibility is further supported by our observation that Osa negatively regulates *Dl* (**Fig 6**). We note that prior studies describe a role of Osa in activation of *Dl* in wing imaginal discs, which contrasts with our observations (Terriente-Félix *et al*. 2009). We attribute this discrepancy as being due to the different spatial patterns and timing of the GAL4 drivers used. Altogether, our data support a direct role for the BAP nucleosome remodeling complex in mediating the proper levels of Notch pathway signaling during wing development.

## METHODS

### Plasmid construction

The *br^disc^*-*tdTomato-PEST* vector was made by cloning a PEST degradation tag from *w; 20xUAS-FLPG5.PEST^attP40^;* (Bloomington 55806), using previously published primers (Nern *et al*. 2011). The PEST sequence was inserted into the previously described pDEST-attR1/2-tdTomato by HiFi assembly (Uyehara *et al*. 2017). The *br^disc^*enhancer was moved into the destination vector by Gateway cloning (Invitrogen). The reporter was integrated into the *attP2, VK33,* and *86FB* landing sites. The *br^disc^-FRT-tdTomato-2xSTOP-FRT-myrGFP* (*br^disc^-switch*) reporter was generated from pJFRC177 *10xUAS-FRT-2xSTOP-FRT-myrGFP* (Addgene 32149). The *br^disc^* enhancer was restriction cloned into the *HindIII* and *AatII* sites, replacing the upstream UAS elements. *TdTomato* cDNA sequence was subsequently restriction cloned into the *NheI* site. The reporter was integrated into the *attP2* landing site. Genomic insertions were made via PhiC31 integration. Injections were performed by BestGene.

Primers used:

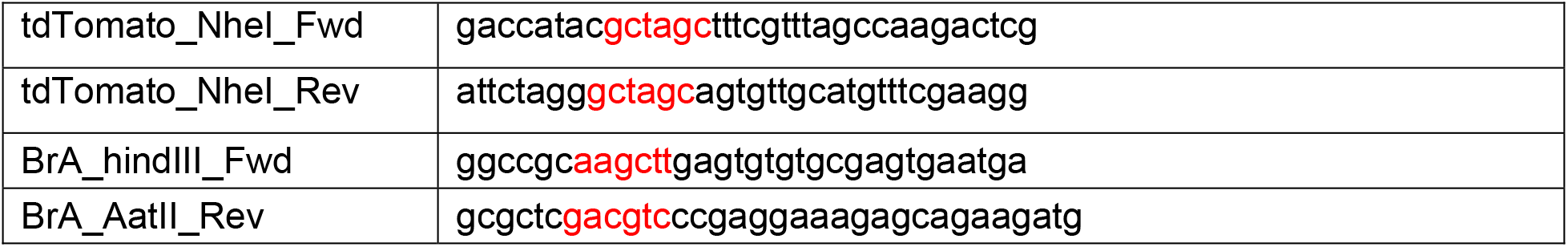
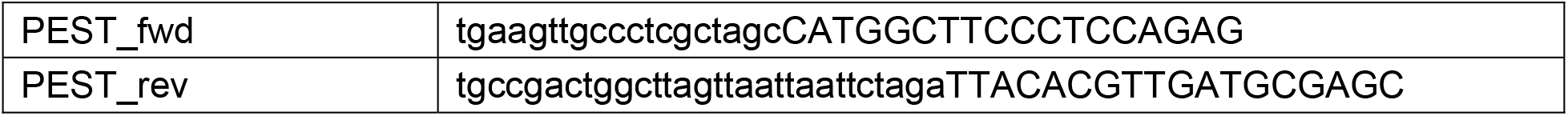

### *Drosophila* culture and genetics

For *br^disc^*-*switch* experiments RNAi expression was driven by *en-GAL4, tub-GAL80^ts.10^* driver. Crosses were raised at 23°C until being shifted to 29°C to induce RNAi. Animals were heat shocked at 37°C for 1 hour to induce Flippase (FLP) expression under the control of the heat-inducible *hsFLP* promoter, and then recovered at 29°C for several hours to allow expression of myr-GFP before dissection. For imaginal discs, crosses were moved to 29°C 72-96h after egg laying (AEL), third larval wandering (3LW) animals were heat shocked 48h later, and then recovered for 4 hours before dissection. For pupal wings, crosses were moved to 29°C 96-120h AEL, prepupae (0-12h APF) were staged using the absence of head-eversion as a developmental marker, heat shocked 24 hours later (24-36h APF), and then recovered for either 4 hours (28-40h APF) or 6 hours (30-42h APF).

For RNAi screening using the *br^disc^-tdTomato-PEST^86Fb^* reporter, RNAi expression was driven by *ci-GAL4, UAS-GFP, tub-GAL80^ts.10^*driver. Crosses were raised at 23°C until being shifted to 29°C to induce RNAi at 72-96h or 96-120h AEL, depending on the severity of phenotypes with individual RNAi lines. Prepupae were staged as described above and then dissected 24h later (24-36h APF). The same protocol was followed to evaluate enhancer hyperactivation except imaginal discs were dissected at 3LW.

For Osa-deGrad experiments, females of the genotype *UAS-Nslmb-vhhGFP4*; *osa^GFP^* / *(TM6B, Tb)* were crossed to males with either *nub-GAL4, tub-GAL80^ts.10^*/ *CyO, Tb-RFP*; *osa^GFP^* / *(TM6B, Tb)* or *osa^GFP^*/ *(TM6B, Tb)* for the negative control lacking GAL4. Crosses were raised at 23°C until 3LW stage. Larvae were moved to 29°C, prepupae were staged 12h later, and non-Tubby female pupal wings were dissected 24h later (24-36h APF). For late Osa-deGrad immunofluorescence experiments, females with *en-GAL4, tub-GAL80^ts.10^*; *osa^GFP^* / *TM6B, Tb* were crossed to males with *UAS-Nslmb-vhhGFP4*; *br^disc^-tdTomato-PEST^VK33^, osa^GFP^* / *TM6B, Tb*. Crosses were kept at 23°C until prepupal stage (0-12h APF) and then moved to 29°C to induce degradation. Non-tubby pupae (*osa^GFP^* homozygous) were dissected 30h later (30-42h APF). Tubby pupae were (*osa^GFP^* heterozygous) were used as a negative control. Younger wings (∼28-38h APF) were identified within the staged range of 28-40h APF and by morphology (small size, absence of folding, absence of elongated bristle shafts along the margin). Older wings (>40h APF) were identified within the staged range of 30-42h APF and by morphology (presence of folds, flattened/expanded cells in wing blade, presence of elongated bristle shafts along the margin) (Sobala and Adler 2016; Diaz and Thompson 2017; Guild *et al*. 2005; Choo *et al*. 2020).

For *osa^308^* mitotic clone experiments, males with the genotype *yw122;; br^disc^-tdTomato-PEST ^attP2^, FRT82B, ubi-GFP / TM6B, Tb* were crossed to females with the genotype *yw;; FRT82B, osa^308^ / TM6B, Tb* at 23°C (day 0). On day 5, vials with larvae were heat shocked in a 37°C water bath for 20 minutes, and then recovered at 25°C for 48h. 0-12h APF prepupae were staged (pre head-eversion) and aged for ∼28h before dissection. Wings were stained with mouse anti-Osa (1:200) and goat anti-mouse Alexa 633 (1:1000).

For CUT&RUN, cultures were raised at 25°C.

Lines used:

*yw; en-GAL4, tub-GAL80^ts.10^*; *br^disc^-FRT-tdTomato-2xSTOP-FRT-myr-GFP ^attP2^* / *(TM6B, Tb)*

*yw; ci-GAL4, UAS-GFP, tub-GAL80^ts.10^* / *(CyO)*; *br^disc^-tdTomato-PEST ^86Fb^* / *(TM6B, Tb)*

*yw; ci-GAL4, UAS-GFP, tub-GAL80^ts.10^* / *(CyO)*; *br^disc^-tdTomato-PEST ^VK33^* / *(TM6B, Tb)*

*yw; en-GAL4, tub-GAL80^ts.10^*; *osa^GFP^* / *(TM6B, Tb)*

*yw; UAS-Nslmb-vhhGFP4*; *osa^GFP^*/ *(TM6B, Tb)*

*w; nub-GAL4^AC-62^, tub-GAL80^ts.10^* / *CyO, Tb-RFP*; *osa^GFP^* / *(TM6B, Tb)*

*yw122;; UAS-osa-RNAi ^attP2^* / *(TM6B)* – (Derived Bloomington 31266)

*yw122;; UAS-lexA-RNAi ^attP2^* / *(TM6B) –* (Derived Bloomington 67945)

*yw;; osa^GFP^ / (TM6B, Tb)*

*yw122;; br^disc^-tdTomato-PEST ^attP2^, FRT82B, ubi-GFP / TM6B, Tb*

*yw;; FRT82B, osa^308^ / TM6B, Tb*

*y, sc, v; UAS-Eip93F-RNAi ^attP40^* – (Bloomington 57868; TRiP.HMC04773)

*yw;;*

See **Table S1** for complete list of RNAi lines.

### Immunofluorescence and image analysis

Larvae and pupae were dissected as previously described (Uyehara *et al*. 2017). Primary antibodies: 1:100 mouse anti-Osa (DSHB), 1:100 rat anti-Elav (DSHB), and 1:10 mouse anti-Delta (DSHB). Secondary antibodies: goat anti-mouse Alexa-633 and goat anti-rat Cy5 were used at 1:1000 (Invitrogen). Tissue was mounted in VECTASHIELD (Vector Labs) with 1.5 coverslips.

For image quantification of RNAi screen microscopy, a custom python script was used to compare reporter signal in RNAi expressing versus WT cells in each wing. Briefly, 10-20 slice z-stacks were converted to maximum intensity projections (MIP). Masks were generated of DAPI, GFP-positive RNAi expressing, and DAPI – GFP (GFP-negative) non-RNAi expressing regions. A ratio of mean grey value in GFP-positive and GFP-negative regions was calculated for each wing.

For image quantification of *br^disc^* hyperactivation and Delta immunofluorescence experiments, MIP were made for each wing, and then regions were selected in RNAi-expressing and WT control cells of the imaginal disc pouch for each wing. For *br^disc^* hyperactivation, square regions were selected that straddled the margin. For Delta quantification, regions were manually drawn around the margin from the anterior-posterior boundary (A/P) to the approximate edge of the most distal provein, L2 in anterior and L5 in the posterior. For both experiments mean grey values were measured using ImageJ (Schindelin *et al*. 2012), and a ratio of mean grey value in RNAi versus WT control was calculated. Student’s two-sample *t*-tests were performed in R to calculate significance.

### High throughput sequencing & data analysis

For FAIRE-seq, wings of female pupae were prepared as previously described (Uyehara *et al*. 2017; Uyehara *et al*. 2019). 40 wings were used per biological replicate. Libraries were prepared using the Takara ThruPLEX DNA-seq kit with unique dual-indexes following manufacturer’s specifications and sequenced on an Illumina NextSeq 2000. Adapters were trimmed from paired-end reads using BBmap BBDuk (v38.71), and then aligned to the dm6 *Drosophila* genome assembly with Bowtie2 (v2.3.4.1; Langmead and Salzberg 2012) with the following parameters: ––very-sensitive ––no-unal ––no-mixed ––no-discordant ––phred33 –I 10 –X 700. Aligned reads were filtered using an exclusion list for dm6 from ENCODE (Amemiya *et al*. 2019), and quality filtered (q > 5) with Samtools (v1.9; Danecek *et al*. 2021), and duplicate reads were removed with Picard (v2.2.4). Coverage files were generated with deepTools (v2.4.1; Ramirez *et al*. 2016) and normalized to 1x genomic coverage (RPGC). Peaks were called with MACS2 (v2.1.2; Zhang *et al*. 2008) using standard parameters. z-normalized coverage files were generated with a custom R script (4.1.3) from RPGC normalized files. For visualization, biological replicates were pooled using Samtools (v1.9). Differential peak analysis was performed in R using DiffBind (v3.8.4; Stark and Brown 2011) and DEseq2 (v1.38.3; Love *et al*. 2014). For assignment of “Osa-dependent” peaks into “Increasing,” “Decreasing,” or “Static” categories, each peak was annotated with z-normalized WT FAIRE-seq data from 3LW, 6h, 18h, 24h, 36h, and 44h APF wings (Uyehara *et al*. 2017). A log_2_ ratio was calculated at each timepoint relative to 3LW. “Increasing” peaks were those that had a log_2_FoldChange >= 1 at 24h, 36h, or 44h APF. “Static” peaks were those that had a log_2_FoldChange between –1 and 1 at 24h, 36h, and 44h APF. “Decreasing” peaks were all remaining peaks. Later pupal stages (24h, 36h, 44h) were used for categorization because they corresponded with approximate stage of wings used for Osa-deGrad FAIRE-seq. Pearson correlation heatmaps of z-normalized coverage files were generated using deepTools (3.5.1).

For WT FAIRE-seq timecourse previously published raw data was accessed from GEO GSE131981 (Ma *et al*. 2019). Data was aligned and processed as described above except alignment was run using Bowtie2 with the ––very-sensitive parameter and no additional changes.

For Osa-GFP CUT&RUN female wing imaginal discs from either *yw;;osa^GFP^*or *yw* negative control animals were dissected and processed as previously described (Uyehara *et al*. 2019). 20-22 wing discs were used for each replicate, with a rabbit anti-GFP (1:100, Rockland 600-401-2156), a pAG-MNase (1:100; UNC core; Salzler *et al*. 2023), and 0.5ng of yeast genomic DNA spike-in (gift of Steve Henikoff). Libraries were prepared from the “supernatant” fraction using the Takara ThruPLEX DNA-seq kit with unique dual-indexes and following the manufacturer’s specifications but with a modified amplification step as previously described (Uyehara *et al*. 2019). Libraries were pooled and sequenced on an Illumina NextSeq 2000 with a 75bp read length. Adapters were trimmed from paired-end reads using BBmap BBDuk (v38.71), and then aligned to the dm6 *Drosophila* genome assembly with Bowtie2 (v2.3.4.1) with the following parameters: ––local ––very-sensitive-local ––no-unal ––no-mixed ––no-discordant –– phred33 –I 10 –X 700. Aligned reads were filtered using a custom exclusion list generated from the “supernatant” of IgG negative controls, as well as anti-Flag and anti-GFP experiments in genotypes that lacked either the Flag or GFP epitopes. Peaks shared among all these negative controls were used to make a conservative list of reproducible high-signal regions. This exclusion list included ∼80 regions. Reads were then quality filtered (q > 5) with Samtools (v1.10), and duplicate reads were removed with Picard (v2.2.4). Coverage files were generated with deepTools (v2.4.1) and normalized to 1x genomic coverage (RPGC). Peaks were called with MACS2 (v2.1.2) without a control and using the –nomodel and –nolamda parameters. Z-normalized coverage files were generated with a custom R script (v4.1.3) from RPGC normalized files.

For H3K27ac CUT&RUN, 20 male imaginal wing discs were used per replicate, with a rabbit anti-H3K27ac (1:100, Active Motif #39135). Libraries were prepared from the “pellet” fractions using the Takara ThruPLEX DNA-seq kit as described above. Libraries were pooled and sequenced on an Illumina Novaseq SP with a 75bp read length. Reads were aligned and processed as described above, except peaks were called with standard MACS2 settings and a sheared genomic DNA control.

For CUT&RUN analysis, only Osa-GFP peak calls greater than or equal to the 50^th^ percentile of MACS2 quality scores (qval) and that were identified in both replicates were kept. Osa-peaks were identified as those that passed screens for quality and reproducibility but did not intersect a reproducible control peak. Peak annotation was performed in R using the ChIPseeker package (v1.34.1; Yu *et al*. 2015), and a negative control bootstrapped shuffle of Osa-peaks was generated using the nullranges package (v1.4.0; Mu *et al*. 2023). Peak overlap enrichment analysis for Hairless ChIP and Rotund ChIP-seq (**Fig 5G,J**) was performed in R using ChIPseeker. Osa-peaks were clustered by dynamic accessibility patterns (**Fig 5H**) by annotating peaks with replicate pooled and z-normalized WT FAIRE-seq data at 3LW, 6h, 18h, 24h, 36h, and 44h APF. For each peak, the fraction of max FAIRE signal was calculated for each timepoint. K-means clustering was performed in R with a k of 8, based on previously described 8 distinct clusters of FAIRE patterns using this data (Nystrom *et al*. 2020). Motif enrichment analysis was performed in R using the memes package (v1.6.0; Nystrom and McKay 2021) and the AME software (McLeay *et al*. 2010; Nystrom *et al*. 2021).

For Rotund ChIP-seq, raw sequencing data was downloaded from the Gene Expression Omnibus (GEO) database (GSE203208, Loker *et al*. 2022). Rotund ChIP-seq data was processed using snakePipes (v2.7.3, Bhardwaj *et al*. 2019). Reads were aligned to dm6 with Bowtie2 (v2.4.5), and peaks were called with MACS2 (v2.2.7.1).

For Hairless ChIP-chip analysis, peak calls were downloaded from GEO (GSE97603, Chan 2017).

All plots were generated in R using the ggplot2 package (v3.4.2), and genome browser plots were generated with the Gviz package (v1.42.1)

### Data Availability Statement

Strains and plasmids are available upon request. High-throughput sequencing data is publicly available online at GEO. Code used to process sequencing data and generate plots can be found at https://github.com/mniederhuber/Niederhuber_2023.

## Figure Legends

**Table S1.** Table of lines used in the RNAi screen. “OTE” denotes any predicted non-specific Off Targets. “Lethality” is a qualitative measure of pupal death in each RNAi experiment. “High” lethality denotes most or all animals with RNAi expression die before eclosion. “Low” lethality denotes some animals die before eclosion, but most survive. “None” lethality denotes most or all animals expressing RNAi survive to adulthood. “Wing phenotype” provides descriptions of any observed wing phenotypes. “KD/WT” mean and standard deviation (SD) columns list quantification of changes in enhancer activity following RNAi expression as plotted in **Fig 2D**. “n” describes the total number of unique wings imaged and quantified for each RNAi line.

**Figure S1.** (A) Confocal images of *br^disc^-GAL4* reporter driving nascent expression of *UAS-myrGFP* in younger (29h APF) and older (44.5h APF) pupal wings. Yellow arrow highlights late activation of *br^disc^* in the elongating bristles of the anterior margin. Images are maximum intensity projections. Scale bars are 100µm.

**Figure S2.** (A) Confocal images of increased *br^disc^* reporter activity following *E93-KD* in the anterior (top-half, outlined with yellow dashed line) compartment relative to WT cells of the posterior (bottom-half), with or without the addition of the PEST degradation tag. Reporter activity from two independent attP integration sites (attP2 and 86Fb) are shown. (B) Illustration of experimental setup for RNAi screen. (C). Confocal images of reduced *br^disc^*reporter activity following *Iswi* knockdown (line 31111) in the RNAi screen. GFP marks the region of RNAi expression. Yellow arrow highlights reduced reporter activity in RNAi-expressing cells. (D) Confocal images of *br^disc^*reporter activity following *polybromo* knockdown (line 330189). (E) Confocal images of *br^disc^* reporter activity in loss-of-function *osa^308^* mitotic clones with anti-Osa immunofluorescence. *Osa^308^* homozygous clones are GFP-, and wild-type twin spots are strong GFP+. Red dashed line marks area of clone. All images are maximum intensity projections. Scale bars are 100µm unless noted.

**Figure S3.** (A) Illustration of GFP insertion in the *osa^GFP^* allele. (B) Confocal images of Osa-GFP and anti-Osa immunofluorescence in wing imaginal discs following *osa* knockdown (line 31266) with or without the posterior *en-GAL4, tub-GAL80^ts^* (*en^ts^*) driver (right-side). (C) Confocal images of Osa-GFP and anti-Osa immunofluorescence in ∼24-36h APF pupal wings following Osa-deGrad with conditions used for FAIRE-seq, with or without the *nub-GAL4, Tub-GAL80^ts^*(*nub^ts^*) driver. Yellow boxes highlight zoomed regions showing changes in nuclear anti-Osa immunofluorescence signal following Osa-deGrad. (D) Venn diagram of Osa-deGrad and Control FAIRE-seq peak calls. (E) Pearson correlation heatmap of z-normalized Osa-deGrad and Control FAIRE-seq coverage files. Microscopy images are maximum intensity projections. Scale bars are 100µm.

**Figure S4.** (A) Confocal microscopy of *br^disc^* reporter hyperactivation in imaginal wing discs following target gene knockdown with an independent *osa* RNAi line (line 330350) and *brm* RNA line (line 31712). GFP marks the region of *ci^ts^* and RNAi expression (yellow dashed line). (B) Confocal microscopy of *br^disc^* reporter hyperactivation and anti-Br immunofluorescence following *osa* knockdown (line 31266) with the *ci^ts^* driver. Images are all maximum intensity projections. Scale bars are 100µm.

**Figure S5.** (A-D) Genome browser shots of z-normalized Osa (magenta) CUT&RUN, Control (grey) CUT&RUN, H3K27ac CUT&RUN (black), and WT FAIRE-seq (black) in wing imaginal discs. Tracks are annotated with Osa peaks (magenta bars and highlights), and Hairless ChIP peaks (teal bars). (E) Heatmap of k-means clustered Fraction of Max z-normalized WT FAIRE-seq signal within distal Osa peaks (see **Methods**). (F) Stacked barplot of Osa peaks that do not overlap a FAIRE peak in imaginal wing discs (3LW) grouped by if those peaks overlap any FAIRE peak at a later stage in the WT FAIRE-seq timecourse.

**Figure S6.** (A) Confocal images of *br^disc^-switch* reporter activity and anti-Elav immunofluorescence in ∼40h APF pupal wings following either *osa* knockdown (line 31266) or control *lexA* knockdown (line 67945) with the posterior (bottom-half) *en^ts^* driver. Zoom inset shows ectopic bristles with nascent myr-GFP signal near to with Elav+ nuclei. All images are maximum intensity projections. Scale bars are 100µm.

